# Decoding Neural Patterns for the Processing of Fearful Faces under Different Visual Awareness Conditions: A Multivariate Pattern Analysis

**DOI:** 10.1101/2022.12.19.520904

**Authors:** Zeguo Qiu, Xuqian Li, Alan J. Pegna

## Abstract

Mixed findings have been reported for the nonconscious processing of fearful faces. Here, we used multivariate pattern analysis on electroencephalography data from three backward masking experiments to decode the conscious and nonconscious processing of fearful faces. Three groups of participants were shown pairs of faces that were presented either subliminally (16 ms) or supraliminally (266 ms) and were required to complete tasks where the face stimuli were either task-relevant (Experiment 1) or task-irrelevant (Experiments 2 and 3). We decoded the neural activity to examine the temporal dynamics of visual awareness, and to investigate whether the presence and location of a fearful face were processed when levels of awareness varied. The results reveal that the spatial location of fearful faces can be decoded from neural patterns only when they are consciously seen and relevant to participants’ task. Nevertheless, the processing of the mere presence of fearful faces can occur in the absence of visual awareness, and the related neural patterns can be generalised to the conscious, non-spatial processing of fearful faces. Additionally, the flexibility of spatial attention seems to modulate the processing of fearful faces.

## Introduction

Negative emotions, in particular fearful expressions, are usually indicative of potentially dangerous events in our surroundings. Our attention is therefore easily attracted, under some circumstances involuntarily, by fearful faces. Observations of the processing of emotional expressions like fear in the absence of awareness are common in the literature. In healthy individuals, the nonconscious processing of emotional faces is usually studied using experimental techniques to suppress normal viewings of the face stimuli (e.g., masking techniques), or by diverting participants’ attention away from the emotional information about the faces (e.g., inattentional blindness; for reviews please see Qiu et al., 2022a; Schindler & Bublatzky, 2020; Tao et al., 2021). It has been consistently shown that even in the unaware conditions, participants could distinguish emotional from neutral expressions, and functional imaging data has shown that several cortical and subcortical regions are activated more strongly by emotional faces in unaware conditions, compared to neutral faces (for a review see Qiu et al., 2022a).

Convergent evidence has been provided from studies on patients with cortical blindness. Patients who experience blindness due to damages to visual cortices have demonstrated preserved detection of emotional expressions despite the impossibility of consciously perceiving visual stimuli, a phenomenon termed “affective blindsight” (Ajina et al., 2010; Burra et al., 2019; De Gelder et al., 1999; Pegna et al., 2005; Striemer et al., 2019). Similarly, hemianopic patients with lesions in their central visual pathway have demonstrated capability of processing emotional information presented in their blind visual field (De Gelder et al., 2005; for a review see Làdavas & Bertini, 2021). These observations are taken to show that, without a functioning visual pathway that enables conscious visual processing, emotional information such as emotional faces can still be processed.

In healthy individuals, backward masking is one of the most common techniques used to manipulate stimulus visibility and hence visual awareness. By presenting target stimuli very briefly (e.g., 16 ms) and presenting backward maskers immediately afterwards, the normal visual processing necessary for awareness in the primary visual cortex is thought to be disrupted (Macknik & Livingstone, 1998). In a series of experiments, we presented face pairs either subliminally (16 ms) or supraliminally (266 ms) and examined the electrical signals, specifically event-related potentials (ERPs), for visual awareness and spatial attention shifting towards a target stimulus (Qiu et al., 2022b, 2022c).

We showed that awareness correlated with two ERPs at an early and a later stage of visual processing, respectively: the Visual Awareness Negativity (VAN) at posterior electrode sites and the P3 or Late Positivity (LP) over parietal electrode sites. However, under the univariate ERP analysis framework, it was impossible for us to examine the temporal dynamics of awareness. Specifically, are there common neural patterns at these two distinct stages of awareness?

Additionally, we found that, the ERP indicative of spatial attention shifting to a lateralised fearful face, indexed by the N2-posterior-contralateral (N2pc), was present only in the supraliminal viewing condition when participants had to localise the fearful face (hence faces were task-relevant; Qiu et al., 2022b). However, no N2pc was found for fearful faces in subliminal conditions, or when the faces were task-irrelevant (Qiu et al., 2022b, 2022c). We concluded that spatial attention shifting towards fearful faces requires visual awareness and is contingent on task-relevancy of the faces (Qiu et al., 2022b, 2022c). At first glance, these results seem to contradict the literature on nonconscious emotion processing. We argue that while the spatial location of fearful faces was not processed nonconsciously, it cannot be inferred that the processing of the faces did not occur at all during subliminal viewings.

Indeed, we found ERP evidence for the (non-spatial) processing of fearful faces in a subliminal viewing condition. Specifically, in a task where participants were asked to localise a lateralised contrast-induced tilted line imposed onto the face images while ignoring the overlapping face stimuli, the ERPs over posterior electrode sites in the fearful-face-present trials were more negative than those in the fearful-face-absent trials (Qiu et al., 2022c). We interpreted this as an overall enhancement in the electrical signals associated with the mere presence of a fearful face, as the manifested effect was not related to the spatial location of the fearful face (Qiu et al., 2022c). It therefore seems that, while the processing of spatial information about the faces requires awareness, the mere presence of fearful faces may be processed in the brain without conscious awareness.

The nonconscious processing of emotional faces has been suggested to function in a coarse and non-specific manner. Detailed or specific information about the faces such as the location of a fearful face may not be encoded without awareness. According to the recurrent processing model (Lamme, 2003), the level of feedback or recurrent activity between neurons determines the fate of visual information. Specifically, subliminally presented stimuli can elicit the fast feedforward activity in the visual cortex, but they fail to initiate the subsequent recurrent activity that is necessary for awareness. While it is possible that stronger activation can be evoked by a subliminally-presented fearful face compared to a neutral face at the initial feedforward stage (e.g., Pessoa & Adolphs, 2010, 2011), the differences in the neural activation may not be sufficient for further processing of other aspects of the stimulus such as its spatial location. Our previous ERP results are consistent with this by showing that the spatial location of a target fearful face can be processed only in the aware conditions (Qiu et al., 2022b), yet a fearful face nevertheless enhances early posterior ERPs regardless of its spatial location when awareness is impeded (Qiu et al., 2022c).

Our next question is whether any nonconscious processing of a fearful face is functionally similar to the conscious processing. Previous research has suggested that nonconscious processing of emotional information is mainly enabled by subcortical pathways (LeDoux, 2000). Specifically, in lesioned brains where the visual cortices are damaged, the pathway through which normal visual processing takes place is destroyed. The preserved or residual capability of detecting emotional expressions in these patients must therefore be enabled by alternate neural paths. Therefore, it has been suggested that the pathways for conscious and nonconscious processing of emotion are subserved by different and somewhat specialised neural connections (Làdavas & Bertini, 2021; Williams et al., 2006). However, existing evidence also provides that cortical regions and several subcortical regions including the amygdala are activated in both conscious and nonconscious processing of fear (for a review see Tao et al., 2021). Hence, it remains unclear whether conscious and nonconscious processing of emotional information are similar to each other and what their neural substrates may be.

To address the questions proposed above, in the current study, we used Multivariate Pattern Analysis (MVPA) to examine the brain activity patterns for processing fearful faces under different conditions of awareness using three datasets from our previous backward masking experiments (Qiu et al., 2022b, 2022c). Electroencephalography (EEG) was recorded while three separate groups of participants performed a fearful face localisation task in Experiment 1, a line comparison task in Experiment 2 and a tilted line localisation task in Experiment 3. With MVPA, signals from each voxel/electrode are not treated independently as in conventional univariate analyses. Instead, the MVPA considers the relationships between multiple voxels/electrodes and “decode” intended experimental manipulations from patterns of brain response distributed across cortical regions (Grootswagers et al., 2017; Haxby et al., 2014). Above-chance decoding gives us confidence to infer that the neural patterns contain information that distinguishes between the experimental conditions. Applying MVPA to EEG data further allows us to examine the temporal dynamics of such decoding (Grootswagers et al., 2017).

Our first aim was to use MVPA to decode awareness and compare the results to our previous ERP findings. Then, we aimed to examine the multidimensional neural patterns during the processing of fearful faces by decoding both the mere presence of a fearful face and its spatial location, under different awareness conditions. Finally, we aimed to examine whether the decoding of fearful faces is generalisable across aware and unaware conditions.

## Methods

### Data description

We used data from our previously published work, consisting of three experiments. The sample sizes were 30, 26 and 20 respectively. The face stimuli were obtained from the Radboud Faces Database (Langner et al., 2010). All materials and apparatus used were the same across three experiments (for a detailed description please see Qiu et al., 2022b, 2022c). Here briefly, we presented face pairs bilaterally to the participants. Three face combinations were presented equiprobably in all experiments: (a) Fearful-Left (where a fearful face was presented to the left of a central fixation with a neutral face to the right; Fig. 1a); (b) Fearful-Right (where a fearful face was presented to the right with a neutral face to the left; Fig. 1b); and (c) Neutral-Control (where two neutral faces were presented; Fig. 1c).

**Fig. 1.**
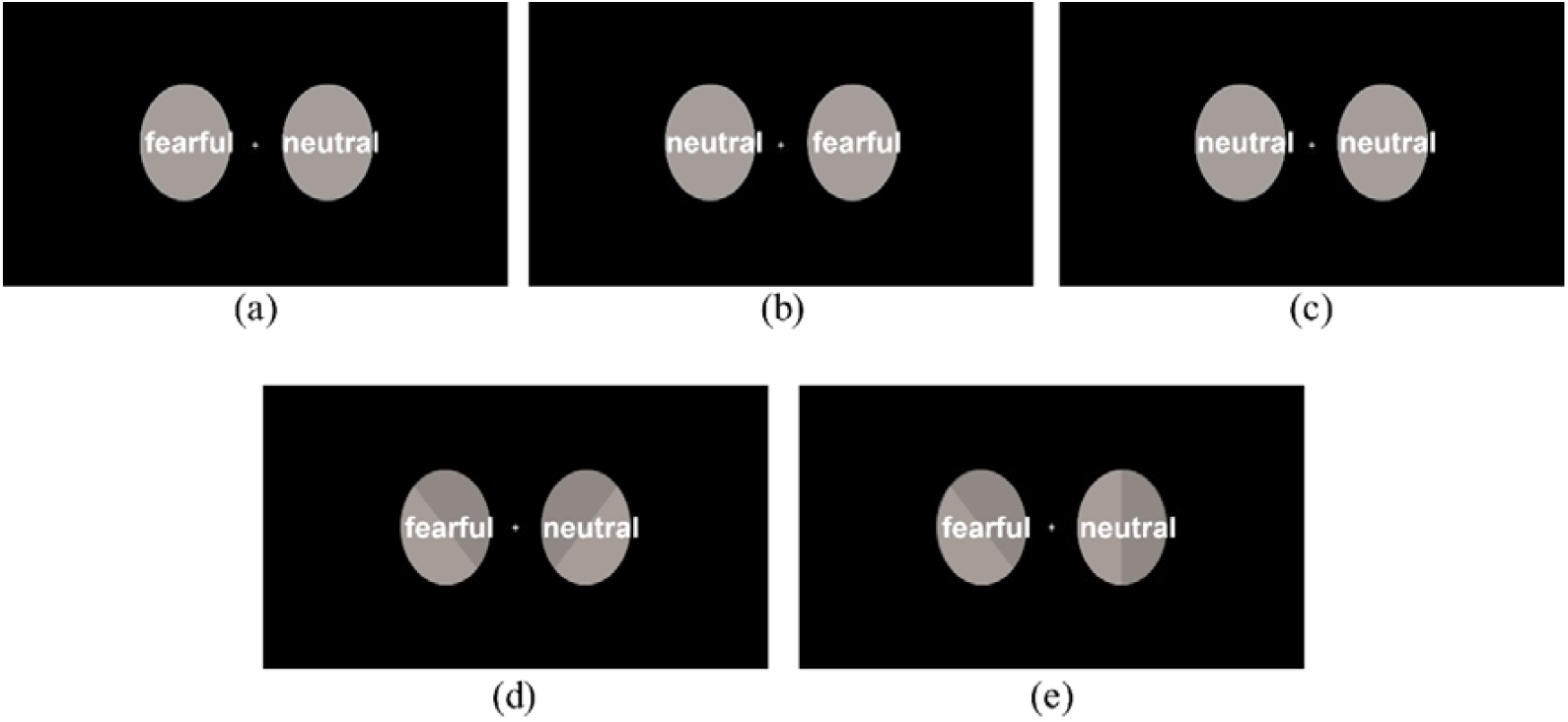
Examples of the stimuli: (a) Fearful-Left, Neutral-Right; (b) Fearful-Right, Neutral-Left; (c) Neutral-Neutral Control; (d) Lines-Different; (e) Tilted-line-on-Left. The face stimuli are covered and de-identified in this picture in accordance with bioRxiv policies. Actual face images were used in the experiment.

In Experiments 2 and 3, a contrast-induced line was additionally imposed onto each face image (see Fig. 1d & 1e). In Experiment 2, each line was either tilted to the left (counter-clockwise) or right (clockwise) such that the bilateral lines were tilted in either a same direction or different directions, see Fig. 1d. In Experiment 3, there were three line combinations: (a) Tilted-line-on-Left (where a tilted line was presented to the left of a central fixation with a vertical line presented to the right); (b) Tilted-line-on-Right (where a tilted line was presented to the right with a vertical line to the left); and (c) Vertical-Control where two vertical lines were presented. The location of the target tilted line and the location of a fearful face were manipulated orthogonally in Experiment 3, see Fig. 1e.

All faces were presented either subliminally (for 16 ms) or supraliminally (for 266 ms) and they were immediately followed by a pair of mask stimuli which were created by randomly scrambling the neutral face image from the same model of the face pair (Fig. 1f). The mask stimuli were presented for 324 ms and 74 ms in the subliminal and the supraliminal conditions, respectively. A blank screen of 550 ms was presented before participants made a behavioural response to the stimuli.

In Experiment 1, participants were required to localise a target fearful face by using the mouse (left click = *Fearful face on the left*; right click = *Fearful face on the right*; middle click = *No fearful face*). In Experiment 2, participants were required to compare the directions two bilateral lines were tilted in by using the mouse (left click = *Same*; right click = *Different*). In Experiment 3, participants were required to localise a target tilted line by using the arrow keys (left arrow key = *Tilted line on the left*; right arrow key = *Tilted line on the right*; down arrow key *= No tilted line*). Detailed descriptions of the experimental procedures can be found in Qiu et al. (2022b, 2022c). While the target was lateralised in both Experiments 1 and 3, there was no lateralised target in Experiment 2 and participants needed to attend to both lines presented bilaterally for the line comparison task.

The faces were task-relevant in Experiment 1 but task-irrelevant in Experiments 2 and 3. In addition, we asked participants to rate their confidence in their responses in Experiments 1 and 2 but not in Experiment 3. For the current study, we did not analyse the data from the task performance or confidence rating.

### EEG recording and pre-processing

Continuous EEG was recorded with a sampling rate of 1024 Hz and the international 10–20 system configuration, using the 64-electrodes BioSemi ActiveTwo system (Biosemi, Amsterdam, Netherlands). Recordings were online referenced to the CMS/DRL electrodes.

Two bipolar electrodes were used to record horizontal electrooculogram (EOG). An external electrode was placed below participants’ left eye and used in combination with FP1 to record vertical EOG.

Pre-processing of the EEG data was performed with EEGLAB (Delorme & Makeig, 2004) and ERPLAB (Lopez-Calderon & Luck, 2014). We interpolated individual electrodes that produced sustained noise throughout the experiment. Signals were re-sampled to 512 Hz offline, filtered from 0.1 to 30 Hz and re-referenced to the average of all electrodes. A notch filter of 50 Hz was applied to remove line noise. ERP signals were segmented into epochs with a time window of 600 ms from the onset of the faces, relative to a pre-stimulus baseline (−100 to 0 ms). Trials with artefacts of eye blinks and eye movements were automatically detected with a threshold of ±100 µV. Trials with small eye-movement artefacts were manually selected and removed. Trials with other artefacts were detected and removed semiautomatically using a threshold of ±80 µV for automatic detection.

### MVPA analyses

#### Decoding visual awareness

In order to test whether the level of visual awareness was decodable from dynamic brain patterns, we applied MVPA to the timeseries data in Experiments 1-3 using the CoSMoMVPA Toolbox (Oosterhof et al., 2016) and A Library for Support Vector Machines (LIBSVM; Chang & Lin, 2011) in MATLAB. For each participant, a radial kernel support vector machine (SVM) was trained at each time point to find the decision boundary that discriminated between patterns of the subliminal and supraliminal conditions using signals across 16 posterior electrodes (P3, P4, P5, P6, P7, P8, P9, P10, PO3, PO4, PO7, PO8, O1, O2, POz, Oz). The pre-processed data were spatially filtered with surface Laplacian (v1.1 CSD Toolbox; Kayser & Tenke, 2006) and temporally smoothed using a 20 ms Gaussian-weighted running average. To improve signal-to-noise ratio while avoiding the overfitting issue, classification was performed on each time point and its 4 neighbouring time points (Grootswagers et al., 2017). Single-trial data in each experiment were partitioned into 10 chunks and such partition was balanced to ensure that each classification target (i.e., subliminal vs. supraliminal) occurred equally often in each chunk. Classifier at each time point was cross-validated using a leave-one-out procedure, with the classifier being trained one nine chunks of data and tested on a non-overlapping chunk. This procedure was repeated to make sure that every chunk was assigned to the test set once. Decoding accuracies of all iterations were then averaged at each time point for each participant.

To further investigate whether neural patterns that discriminated between subliminal and supraliminal conditions remained stable over time, we performed temporal generalisation analysis for visual awareness in each experiment (King & Dehaene, 2014). Instead of being trained and tested at the same time point, a classifier was trained at one time point and tested at all other time points to examine whether neural patterns that was identified at a specific time could be generalised to another time.

#### Decoding the presence of fearful faces

In Experiments 1-3, we also performed decoding analysis at each level of visual awareness to see whether brain patterns related to the mere presence of fearful faces (i.e., Fearful-Left and Fearful-Right) were separable from those of the absence of fearful faces (i.e., Neutral-Control). These analyses followed the same procedures described above except for one additional step. Because all experiments involved a lateralised presentation of a fearful face in the fearful-face-present trials, we flipped data between hemispheres for all Fearful Right conditions, such that in both the Fearful Left and Fearful Right conditions, electrodes in the left hemisphere represented brain activity ipsilateral to the fearful face while electrodes in the right hemisphere represented brain activity contralateral to the fearful face. This was done to maximally capture any lateralised effect by retaining consistency across all trials of the same classification target (i.e., presence). To test whether the decoding of fearful face representations at different levels of visual awareness was driven by any shared neural patterns, successful decoding of the presence of fearful faces was further explored using a cross-condition validation scheme. This procedure allowed us to train classifiers on the subliminal conditions and test on the supraliminal conditions, or vice versa.

#### Decoding the location of fearful faces

Finally, we were interested in the effect of spatial attention, and more specifically, the interaction between visual awareness and spatial attention. We therefore performed decoding analysis in Experiments 1-3 to distinguish between the Fearful Left and Fearful Right conditions at each level of stimulus visibility.

The above-mentioned analyses were tested for statistical significance using one-sample *t*-tests along with the Monte Carlo permutations at the group level to determine whether the observed accuracies were significantly above chance (i.e., 50%). The *t* statistics were corrected for multiple comparisons by computing the threshold-free cluster enhancement (TFCE) statistics (Smith & Nichols, 2009). Null distribution was acquired over 10,000 iterations of randomly flipping the sign of the statistic values across time points. The observed TFCE statistic at each time point was considered significant if its value was larger than the 95th percentile of the null distribution, corresponding to *p* < .05 for a one-tailed test, corrected for multiple comparisons.

### Supplementary analyses

We repeated all the above analyses at the whole-brain level by including all scalp electrodes in the analyses. The effects reported above were fully replicated. A full report of the whole-brain analyses can be found in the Supplementary Materials.

## Results

### The decoding for visual awareness

Neural patterns across posterior brain regions track visual awareness in all three experiments. In Experiment 1 (Fig. 2a), awareness level (subliminal vs. supraliminal) was first decodable at 78 ms after stimulus onset, followed by rapid increases in decoding accuracy to 65% (*SEM* = 1.21) at 158 ms and 71% (*SEM* = 1.42) at 235 ms. The levels of visual awareness continued to be decodable for the rest of the trial, with a third peak at around 445 ms reaching 71% (*SEM* = 1.23). In Experiment 2 (Fig. 2b), the subliminal and supraliminal conditions were reliably distinguishable throughout the trial since 104 ms. The time course of decoding accuracy revealed three major peaks that looked similar to those found in Experiment 1. The first peak was found at 165 ms at 63% (*SEM* = 1.55), followed by the highest peak at 260 ms at 69 % (*SEM* = 1.42) and the third peak at 478 ms at 67% (*SEM* = 1.26). In Experiment 3 (Fig. 2c), the above-chance decoding of awareness level first appeared at 128 ms. This was followed by a small peak at 168 ms and a larger peak at 248 ms that reached 64% (*SEM* = 1.51).

**Fig. 2.**
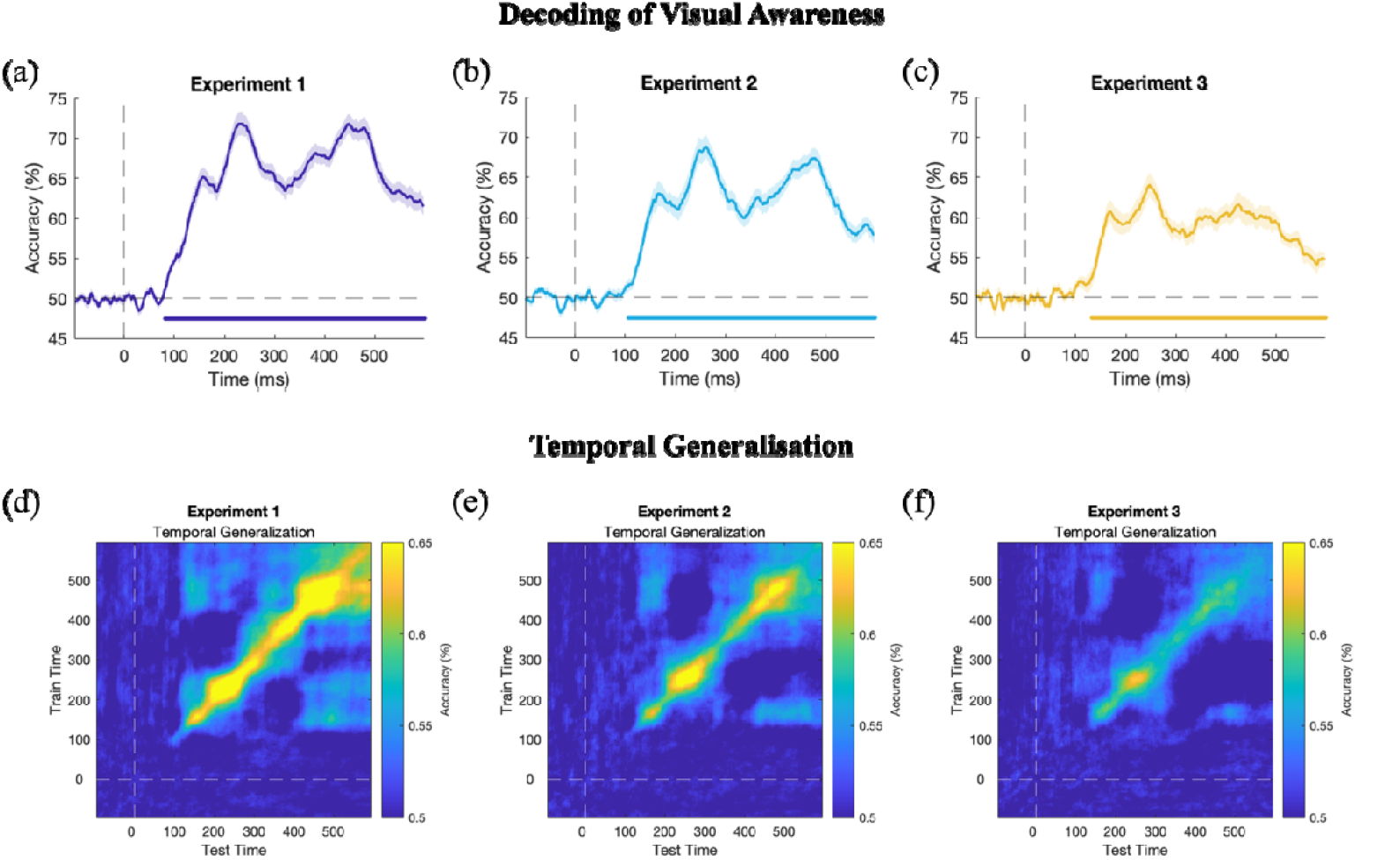
Decoding and Temporal Generalisation of Visual Awareness. Performance of visual awareness decoding (supraliminal vs. subliminal) using EEG signals across posterior electrodes throughout the trial in (a) Experiments 1, (b) Experiment 2, and (c) Experiment 3 are shown. Time periods showing above-chance decoding (accuracy > 50%) are highlighted by the coloured dotted line in each plot, TFCE-corrected *p* < 0.05. The coloured shading in each plot represents standard error of the mean. Temporal generalisation of visual awareness decoding in (d) Experiments 1, (e) Experiment 2, and (f) Experiment 3 are shown.

To test the stability of classifiers throughout the trial, temporal generalisation analysis was carried out for the decoding of visual awareness. In Experiment 1, classifiers from about 110-250 ms were generalisable to a later time period at about 400-600 ms (Fig. 2d); these off-diagonal effects suggested that neural patterns occurring early in the trial that distinguished subliminal from supraliminal viewings may be reactivated later to discriminate the two conditions. A sustained pattern of temporal generalisation was revealed from about 350 ms to the end of the trial; this square-shaped decoding performance may suggest a unitary mechanism that happened late in the trial. In Experiment 2, off-diagonal effects were found when classifiers were trained at around 140-190 ms and tested at around 430-580 ms (Fig. 2e), which suggests the generalisability of neutral patterns at the early stage of visual awareness.

The squared-shape pattern of temporal generalisation was also found in Experiment 2 from 350 ms till the end of the trial. However, in Experiment 3, the generalisation of neural activity at an early stage to a later stage was no longer significant. The above-chance, sustained pattern of temporal generalisation was found from around 400 ms to the end of the trial (Fig. 2f), for Experiment 3.

### The decoding of the mere presence of fearful faces

In Experiment 1, reliable decoding of supraliminal fearful faces (vs. neutral faces) started immediately after stimulus onset, followed by a peak at 138 ms and a peak at 265 ms with an accuracy of 63% (*SEM* = 1.14) and 62% (*SEM* = 0.89), respectively (Fig. 3a).

**Fig. 3.**
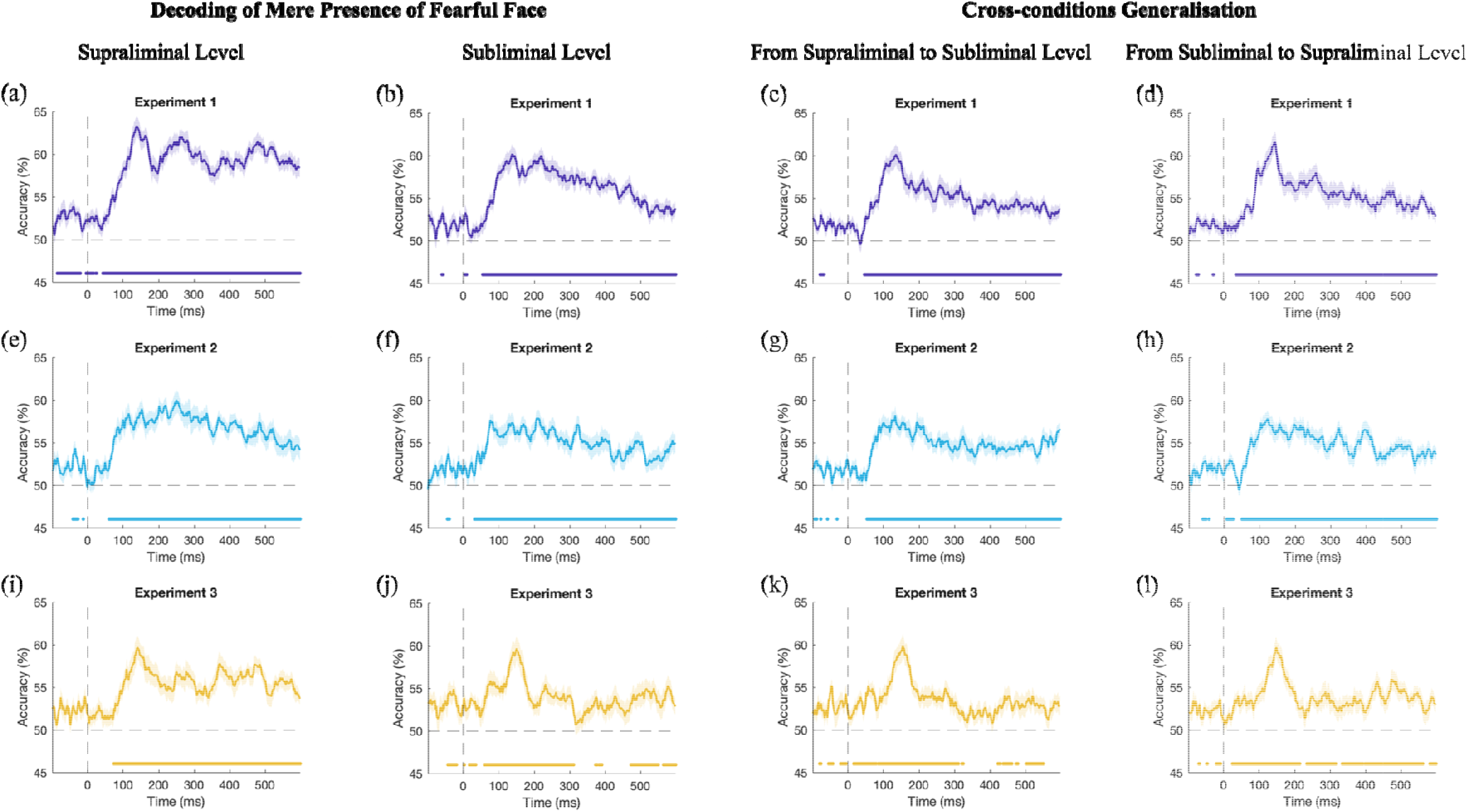
Main Decoding and Cross-condition Decoding of Mere Presence of Fearful Face. Performance of fearful face decoding (presence vs. absence) using EEG signals across posterior electrodes at supraliminal level and subliminal level, and cross-condition decoding of fearful face from supraliminal to subliminal level and from subliminal to supraliminal level, for Experiment 1 (a-d), Experiment 2 (e-h) and Experiment (i-l) are shown. Time periods showing above-chance decoding (accuracy > 50%) are highlighted by the coloured dotted line in each plot, TFCE-corrected *p* < 0.05. The coloured shading in each plot represents standard errors of the mean.

Interestingly, when classifying between subliminal stimuli, above-chance decoding was also found at most of the time points (Fig. 3b). Such decoding first reached a peak at 138 ms with an accuracy of 60% (*SEM* = 1.19) and steadily decreased to 53% (*SEM* = 0.82) at the end of the trial. The cross-condition validation analysis further revealed above-chance decoding when classifiers were trained on the supraliminal conditions and tested on the subliminal conditions, with a peak accuracy of 60% (*SEM* = 1.20) at around 135 ms (Fig. 3c).

Surprisingly, classifiers that were trained on the subliminal conditions also predicted differences between fearful and neutral faces in supraliminal trials, with the decoding accuracy peaking at 62% (*SEM* = 1.22) at 145 ms (Fig. 3d). These findings suggest that neural patterns that distinguished between fearful and neutral faces can be generalised across the two levels of stimulus visibility.

In Experiment 2, above-chance decoding in the supraliminal conditions first appeared at around 57 ms, reached the highest accuracy at 60% (*SEM* = 1.28) at 250 ms, and gradually decreased to around 54% during the rest of the trial (Fig. 3e). Fearful face decoding at subliminal level was significant starting at ∼30 ms post-stimulus; classifiers’ performance remained relatively stable throughout the trial with a mean accuracy of 55% (Fig. 3f). The cross-condition analyses showed significant decoding when subliminal trials were tested on the supraliminal classifiers (Fig. 3g) and when supraliminal trials were tested on the subliminal classifiers (Fig. 3h), with mean accuracies of about 55%. These findings again suggest that brain patterns of the consciously processed fearful faces can be generalised to the nonconscious processing of fearful faces and vice versa.

In Experiment 3, the supraliminal fearful faces were first distinguishable from neutral faces at 70 ms after stimulus onset, with a peak of decoding accuracy at 60% (*SEM* = 1.33) found later at 140 ms (Fig. 3i). Such decoding continued to be above the chance level during the rest of the trial, with the accuracy fluctuating around a mean of 56%. At subliminal level, on the other hand, fearful faces were decodable immediately after stimulus onset, followed by a period of above-chance decoding from 56-315 ms with a peak in accuracy at 60% (*SEM* = 1.38) at 150 ms (Fig. 3j). A relatively sustained period of decoding lasting for 125 ms was also found later from 468 ms to the end of the trial. Similar to Experiments 1 and 2, the cross-condition analysis revealed a generalisability of fearful face classifiers from supraliminal to subliminal conditions and vice versa, with the best decoding performance at around 145-155 ms with accuracies above 59% (Fig. 3k & 3l).

### The decoding for the spatial location of fearful faces

In Experiment 1, the spatial location (left vs. right) of fearful faces were only decodable in the supraliminal conditions (Fig. 4a) but not in the subliminal conditions (Fig. 4b). Successful decoding in the supraliminal conditions was mainly found for time windows of 160-320 ms, 365-510 ms, and 535-595 ms, with a peak at 58% (*SEM* = 0.92) at around 410 ms. A *post-hoc* temporal generalisation analysis suggests that the early classifiers trained at 160-320 ms was not generalisable to later time points (Fig. 4c). In Experiments 2 and 3, no significant decoding of spatial location was found at either awareness level, with mean decoding accuracies ranging from 50% to 51%.

**Fig. 4.**
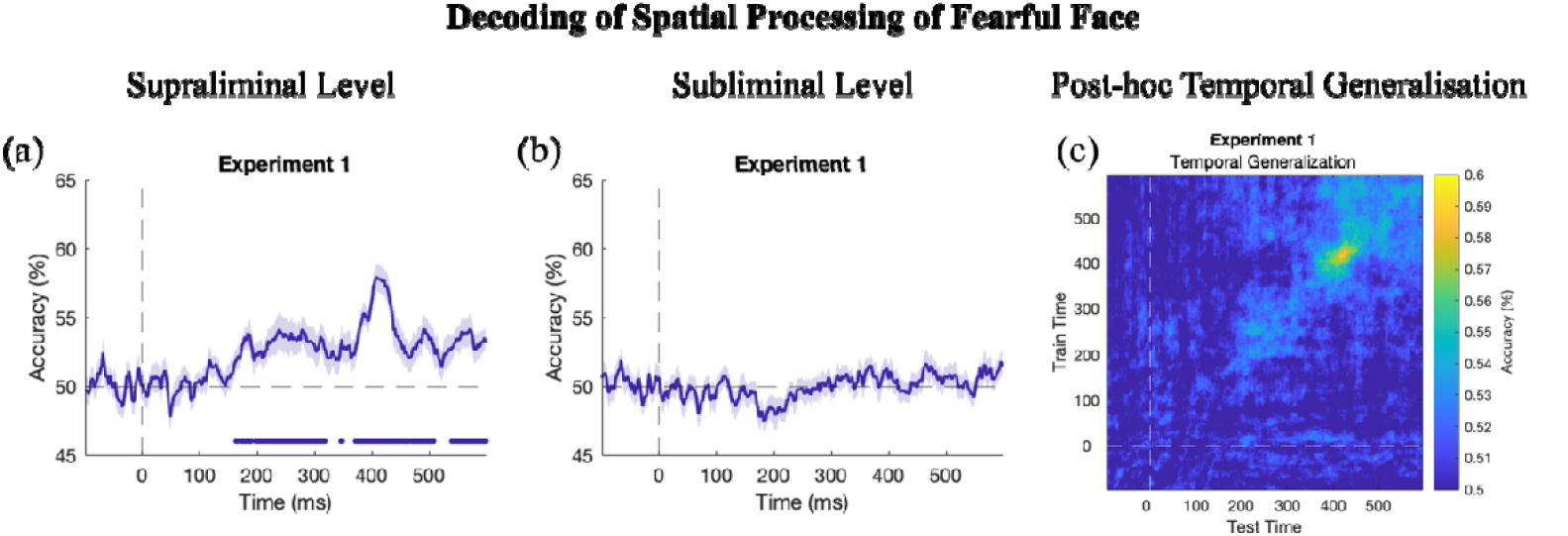
Decoding of Spatial Processing of Fearful Face in Experiment 1. Decoding performance of spatial processing of fearful face (a) at supraliminal level and (b) subliminal level are shown. Time periods showing above-chance decoding (accuracy > 50%) are highlighted by the coloured dotted line in each plot, TFCE-corrected *p* < 0.05. The coloured shading in each plot represents standard error of the mean. (c) Post-hoc temporal generalisation of spatial processing decoding at supraliminal level in Experiment 1.

## Discussion

By performing MVPA on EEG data across three backward masking experiments, we distinguished brain patterns between conditions where pairs of face images were presented subliminally (16 ms) and conditions where they were presented supraliminally (266 ms), with maximal decoding performance observed at three distinct time windows. Additionally, we showed that the presence of a fearful face in the display could be decoded during both subliminal and supraliminal viewings in all three experiments, whereas the spatial location of the fearful face was decodable only in the supraliminal condition and when the faces were task-relevant (in Experiment 1).

### Visual awareness is maximally decodable at three stages

Starting from ∼70 ms in Experiment 1 and ∼100 ms in Experiments 2 and 3, successful decoding of visual awareness was found throughout the duration of the trial. These results showed that information represented in the posterior cortex started to diverge early at around 70 ms post-stimulus between subliminal and supraliminal viewing conditions. The backward masking technique used in the experiments is powerful at suppressing visual awareness due to the interfering visual inputs (i.e., masks; Macknik & Livingstone, 1998). When visual stimuli were presented for 16 ms and immediately backward masked, recurrent neural activity was not initiated sufficiently, if at all (Lamme, 2003). However, in the supraliminal condition, because the stimuli were presented for much longer, ongoing recurrent activity was enabled, leading to an overall enhanced electrical activity in the brain and consequently conscious awareness of the stimuli. Our current results extend beyond previous ERP findings by showing that differences between supra- and subliminal conditions are tracked by different multidimensional neural patterns distributed across the posterior cortex.

Notably, peaks in decoding accuracy were shown in two early time windows in all three experiments: 158-168 ms and 235-260 ms. One additional peak of accuracy appeared after 400 ms for Experiments 1 and 2, and for Experiment 3 but to a lesser extent. These results suggest that posterior electrodes were more sensitive to the effect of stimulus visibility at these distinct stages of processing. The time windows of these accuracy peaks align with the timing of several ERP components of relevance in the awareness and face processing literature (Qiu et al., 2022b, 2022c, 2022d): the face-sensitive N170 (150-180 ms) and two awareness-related components (the VAN at 200-300 ms and the LP at 400-600 ms). We would like to highlight that, prior to the VAN, a component thought to be the first reliable awareness-related electrical marker (Förster et al., 2020), visual awareness of face stimuli can already be robustly decoded at ∼150 ms post-stimulus.

According to Lamme’s recurrent processing account of consciousness (Lamme, 2003, 2010), the emergence of visual awareness results from feedback activity between neurons that is largely localised to early visual areas starting at ∼80 ms. As information processing progresses, such feedback activity becomes more widespread at a large scale of neural network (Lamme, 2010). The earlier time windows with high decoding accuracy (∼150-300 ms) likely correspond to the initial stages of recurrent processes and thus the early stage of visual awareness, while the late peak (after 400 ms) is driven by the wider-spread recurrent activity for a later stage of awareness.

Importantly, our temporal generalisation analysis revealed that the awareness-related classifier at ∼200 ms can be generalised to 400-600 ms in Experiments 1 and 2, indicating that neural patterns that distinguished levels of visual awareness at ∼200 ms reoccurred later after 400 ms in the posterior cortex. Note that the temporal generalisation result was not significant in Experiment 3 whereby the classifier for the supraliminal-subliminal decoding at ∼200 ms did not reoccur in later stages of decoding. This could be due to a low sample size in Experiment 3, especially considering that numerous classification tests were performed in the temporal generalisation analysis (354 × 354 = 125316).

### Processing of the location of fearful faces is contingent on awareness and task-relevancy

We found that the classification between Fearful-Left and Fearful-Right trials was successful, however only in the supraliminal condition when the faces were task-relevant (in Experiment 1). The decoding of the spatial location of the fearful faces was not possible when the faces were task-irrelevant or when they were presented subliminally. These findings support our previous ERP analyses on the data (Qiu et al., 2022b, 2022c). In those studies, the N2pc, an ERP that indexes spatial attention shifting, appeared for target fearful faces only when the stimuli were presented supraliminally. Here, by examining neural patterns via a multivariate technique, we confirm that the processing of the spatial location of faces requires visual awareness.

However, awareness is not sufficient for localising a fearful face spatially, as shown by chance-level performance in the location decoding in Experiments 2 and 3. In these two experiments, the faces were made irrelevant to participants’ task goals. Perhaps, spatial information about the fearful faces were suppressed to allow the task-relevant information (i.e., contrast-induced lines) to be accurately represented in the brain. Consequently, the spatial location of irrelevant fearful faces was not encoded by the brain. The top-down active suppression of task-irrelevant information is an important mechanism in the attentional capture literature (for a review see Luck et al., 2021). Numerous studies have shown that task-irrelevant information, albeit salient (e.g., colour singletons), does not capture attention, possibly because the stimulus does not match participants’ internal attentional template (Luck et al., 2021).

### The mere presence of a fearful face is decoded regardless of visual awareness

Nevertheless, the neural processing of fearful faces was not absent in task-irrelevant settings. Our current results showed that the mere presence of a fearful face was successfully decoded in both subliminal and supraliminal viewing conditions across all three experiments. It therefore appears that the processing of a fearful face in the visual display, regardless of its location, is possible even in the absence of conscious awareness.

Fearful faces have been consistently shown to enhance neural activity than their neutral counterparts and other emotional expressions (for a review see Schindler & Bublatzky, 2020). The prioritised processing of fear is not only observed in normal viewings, but also preserved in situations where cortical responses are impeded by using experimental manipulations (e.g., backward masking), or in circumstances of cortical blindness caused by brain damages (i.e., affective blindsight; De Gelder et al., 1999). It has been suggested that fear-related signals can reach the amygdala, a subcortical region specialised for processing fear, bypassing the visual cortex (e.g., via the colliculus-pulvinar-subcortical pathway; LeDoux, 2000). The existence of such subcortical pathways has been increasingly supported since it was proposed (Compton, 2003; Framorando et al., 2021; Méndez-Bértolo et al., 2016; Tamietto & De Gelder, 2010; for a recent review see Dahlén et al., 2022).

Consistent with these claims, our MVPA results showed that, when visual awareness was impeded during subliminal viewings, the presence of a fearful face in the display nevertheless changed the neural responses across brain regions in healthy adults. While it is likely that the presence of a fearful face can be readily encoded during the initial fast feedforward stage of visual processing (Pessoa & Adolphs, 2010, 2011), we argue that these changes may also be enabled, at least partly, by the subcortical pathways. Possibly, the amygdala is more strongly activated in response to a fearful face and the signals can be subsequently passed on to the occipital areas via direct connections (Bayle et al., 2009; Catani et al., 2003; Vuilleumier, 2005). Supporting this explanation, the timing of the peak in the presence decoding is consistent with the effect related to the amygdala responses to emotional faces (as early as 74 ms in Méndez-Bértolo et al., 2016). Nonetheless, we have to emphasise that this explanation is speculative and future functional imaging studies are needed to test the involvement of the amygdala during the process.

Our results also revealed successful generalisation of the presence decoding between subliminal and supraliminal conditions, suggesting that the differences in neural patterns between Fearful-face-present and Fearful-face-absent trials in the subliminal conditions are somewhat in common with those in the supraliminal conditions, and vice versa. Previous research has found similarity in the activated regions between conscious and nonconscious processing of emotional faces (Fogazzi et al., 2020; Prete et al., 2015; Prochnow et al., 2013; Tao et al., 2021). Perhaps, while the initial stages of visual processes is impeded in the subliminal condition, the information can nevertheless be conveyed to other regions (i.e., higher-level visual cortices and fronto-parietal regions; LeDoux, 2000) via alternate pathways that are less affected by procedures such as masking, as explained above.

Crucially, the successful decoding of the presence but not the spatial location of a fearful face in the subliminal condition suggests that nonconscious processing of fearful faces is not spatially constrained. This is consistent with previous findings from studies on affective blindsight. Specifically, using an attention-shifting paradigm with visual and auditory stimuli, Burra et al. (2014) presented faces with either direct or averted (to the left or right) eye gaze to a blindsight patient with complete destruction of the primary visual cortex. The patient was required to localise a subsequent auditory stimulus presented either to his left-or to his righthand side. Contrary to healthy controls, direction of eye gaze did not affect the patient’s responses to the auditory stimuli, showing that attention was not oriented by gaze cues in the absence of awareness. In an fMRI study, the same patient was presented with faces displaying direct or averted eye gaze (Burra et al., 2013). The activation of the right amygdala was found to be enhanced for presentations of direct eye gaze, compared to averted gaze. These findings were taken to suggest that nonconscious processing of meaningful eyegaze information is possible, however gaze-cued attention orientation is abolished in the absence of visual awareness. Moreover, the subcortical pathway was proposed to be the main neural pathway for the non-spatial nonconscious processing of eye gaze information (Burra et al., 2013, 2014).

### The level of attention flexibility affects the neural encoding of task-irrelevant information

Finally, when visually inspecting the time courses of fearful face presence decoding across three experiments, we found a peak in the decoding accuracy for Experiments 1 and 3 but not for Experiment 2. Specifically, in an early stage of face processing (i.e., 158-168 ms), we saw a large peak in the decoding accuracy in both subliminal and supraliminal conditions in Experiments 1 and 3, suggesting that the presence of a fearful face was more decodable in this time range compared to other time points. However, this decoding accuracy peak was not found in Experiment 2. We propose that the differences were perhaps due to different levels of attention flexibility across experiments, induced by task requirements.

In Experiments 1 and 3, the target stimulus was lateralised and participants were required to localise a target that could appear on either the left or the right side of the screen. Their covert spatial attention was thus expected to be shifted across the screen. However, in Experiment 2, the task was to compare two lines presented bilaterally and attention was required to be maintained on both sides to allow for an accurate neural representation of both lines. Therefore, the flexibility in attention shifting is naturally lower in the line comparison task (in Experiment 2), compared to the other two experiments. Previous studies have shown that task demands can alter the neuronal activation levels and hence affect neural encoding of the same visual information (Harel et al., 2014; Jackson et al., 2017; Roy et al., 2010; Stokes et al., 2013; Yu & Shim, 2017). Specifically, when categorising similar visual information, the neural representation of task-irrelevant features of the stimuli will be significantly weakened, usually indicated by worse decoding performance, compared to task-relevant features (e.g., Jackson et al., 2017). From a different yet related perspective, in Experiment 2, two individual line stimuli needed to be represented equally well in the brain for generating an answer on how they compare to each other, which is in contrast with Experiments 1 and 3 where only one target stimulus in the pair was sufficient for making a response. A stronger suppression was thus necessitated for two task-irrelevant distractors (face images) in Experiment 2.

Therefore, when task requirements varied across experiments, from allowing spatial attention to be shifted (in Experiments 1 and 3) to forcing equal distribution of attention (in Experiment 2), the depth of neural processing of the fearful faces varied, leading to differential neural decoding patterns. In our current context, this was demonstrated as a much higher accuracy at an early stage of face processing (∼150 ms) in decoding the presence of a fearful face in an attentionally flexible situation (Experiments 1 and 3), compared to the less flexible situation (Experiment 2). Future research should aim to systematically vary the level of attention flexibility and examine its effect on the processing of task-irrelevant information.

In conclusion, the MVPA analysis of three previous experimental studies reveals that the spatial location of fearful faces can be encoded only when they are consciously seen and task-relevant. Importantly, fearful faces are nevertheless processed in the absence of visual awareness, and this processing shares some common neural patterns with the conscious non-spatial processing of fearful faces.

## Data and code availability

Data and codes for Experiments 1 and 2 can be found at https://osf.io/p4zks/. Data and codes for Experiment 3 can be found at https://osf.io/dqbkg/.

## Author contributions

**Zeguo Qiu**: Conceptualization; formal analysis; methodology; project administration; writing – original draft; writing – review and editing. **Xuqian Li**: Formal analysis; methodology; visualization, writing – review and editing. **Alan J. Pegna**: writing – review and editing.

## Acknowledgements

Zeguo Qiu and Xuqian Li are supported by Graduate School Scholarships provided by The University of Queensland.

## Conflict of interest

The authors have no conflict of interest to report.

## Notes

### Competing Interest Statement

The authors have declared no competing interest.

