## Supplementary Materials for "Decoding Neural Patterns for the Processing of Fearful Faces under Different Visual Awareness Conditions: A Multivariate Pattern Analysis"

**Supplementary Results**

**Whole-brain decoding for visual awareness**

Neural patterns across whole-brain electrodes track visual awareness in all three experiments. In Experiment 1 (Fig. S1a), awareness level (subliminal vs. supraliminal) was first decodable at 84 ms after stimulus onset, followed by rapid increases in decoding accuracy to 65% (*SEM* = 1.14) at 160 ms and 75% (*SEM* = 1.43) at 235 ms. The levels of visual awareness continued to be decodable for the rest of the trial, with a third, larger peak at around 445 ms reaching 73% (*SEM* = 1.27). In Experiment 2 (Fig. S1b), the subliminal and supraliminal conditions were reliably distinguishable throughout the trial since 104 ms. The time course of decoding accuracy revealed three major peaks that looked similar to those found in Experiment 1. The first peak was found at 165 ms at 63% (*SEM* = 1.57), followed by the highest peak at 255 ms at 71 % (*SEM* = 1.58) and the third peak at 440 ms at 70% (*SEM* = 1.31). In Experiment 3 (Fig. S1c), the above-chance decoding of awareness level first appeared at 117 ms. This was followed by a small peak at 174 ms and a larger peak at 246 ms that reached 66% (*SEM* = 1.85).

To test the stability of classifiers throughout the trial, temporal generalisation analysis was carried out for the decoding of visual awareness. In Experiment 1, classifiers from about 110-250 ms were generalisable to a later time period at about 400-600 ms (Fig. S1d); these off-diagonal effects suggested that neural patterns occurring early in the trial that distinguished subliminal from supraliminal viewings may be reactivated later to discriminate the two conditions. A sustained pattern of temporal generalisation was revealed from about 350 ms to the end of the trial; this square-shaped decoding performance may suggest a unitary mechanism that happened late in the trial. In Experiment 2, off-diagonal effects were found when classifiers were trained at around 150-200 ms and tested at around 400-580 ms (Fig. S1e), which suggests the generalisability of whole-brain patterns at the early stage of visual awareness. The squared-shape pattern of temporal generalisation was also found in Experiment 2 from 350 ms till the end of the trial. However, in Experiment 3, the generalisation of neural activity at an early stage to a later stage was no longer significant. The above-chance, sustained pattern of temporal generalisation was found from around 400 ms to the end of the trial (Fig. S1f), for Experiment 3.

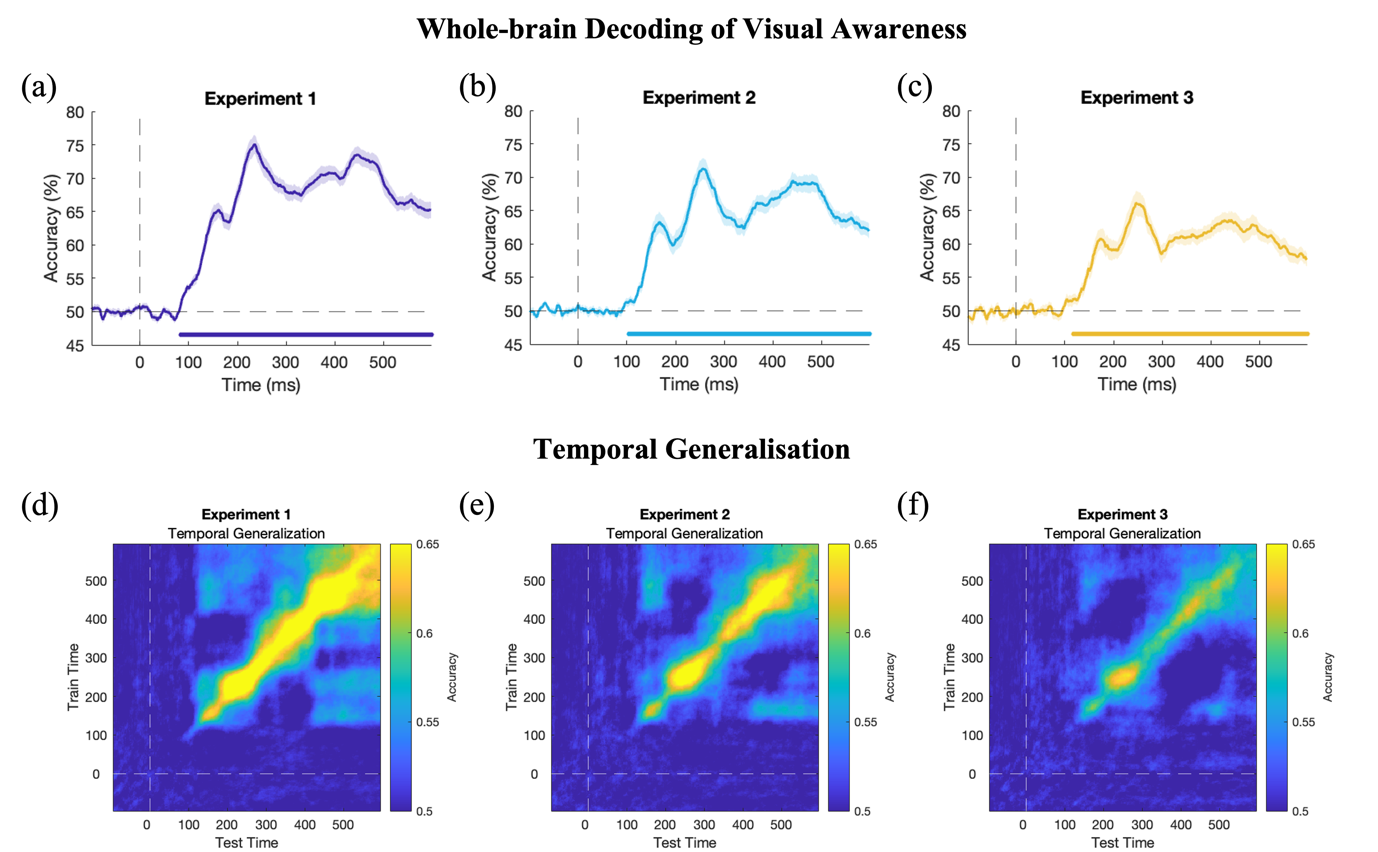

**Fig. S1.** Whole-brain Decoding and Temporal Generalization of Visual Awareness. Performance of visual awareness decoding (supraliminal vs. subliminal) using EEG signals across whole-brain electrodes throughout the trial in (a) Experiments 1, (b) Experiment 2, and (c) Experiment 3 are shown. Time periods showing above-chance decoding (accuracy > 50%) are highlighted by the coloured dotted line in each plot, TFCE-corrected *p* < 0.05. The coloured shading in each plot represents standard error of the mean. Temporal generalization of visual awareness decoding in (d) Experiments 1, (e) Experiment 2, and (f) Experiment 3 are shown.

**Whole-brain decoding of the mere presence of fearful faces**

In Experiment 1, reliable decoding of supraliminal fearful faces (vs. neutral faces) started at 45 ms after stimulus onset, followed by a peak at 143 ms and a peak at about 468 ms with an accuracy of 63% (*SEM* = 0.82) and 63% (*SEM* = 0.99), respectively (Fig. S2a). Interestingly, when classifying between subliminal stimuli, above-chance decoding was also found at most of the time points (Fig. S2d). Such decoding first reached a peak at 142 ms with an accuracy of 60% (*SEM* = 1.09) and steadily decreased to about 55% (*SEM* = 0.71) at the end of the trial. The cross-condition validation analysis further revealed above-chance decoding when classifiers were trained on the supraliminal conditions and tested on the subliminal conditions, with a peak accuracy of 60% (*SEM* = 1.31) at around 138 ms (Fig. S2g). Classifiers that were trained on the subliminal conditions also predicted differences between fearful and neutral faces in supraliminal trials, with the decoding accuracy peaking at 61% (*SEM* = 1.11) at 140 ms (Fig. S2j).

In Experiment 2, above-chance decoding in the supraliminal conditions appeared immediately after stimulus onset, reached accuracy between 57%-59% from 100 to 400 ms, and gradually decreased to around 54% during the rest of the trial (Fig. S2b). Fearful face decoding at subliminal level was significant starting at stimulus onset; classifiers’ performance remained relatively stable throughout the trial with a mean accuracy of 55% (Fig. S2e). The cross-condition analyses showed significant decoding when subliminal trials were tested on the supraliminal classifiers (Fig. S2h) and when supraliminal trials were tested on the subliminal classifiers (Fig. S2k), with mean accuracies of about 54%. These findings again suggest that whole-brain patterns of the consciously processed fearful faces can be generalised to the unconscious processing of fearful faces and vice versa.

In Experiment 3, the supraliminal fearful faces were reliably distinguishable from neutral faces at 72 ms after stimulus onset, with a peak of decoding accuracy at 57% (*SEM* = 1.47) found later at 140 ms (Fig. S2c). Such decoding continued to be above the chance level during the rest of the trial, with a second peak at 57% (*SEM* = 1.03) at 360 ms. At subliminal level, fearful faces were decodable since 62 ms after stimulus onset, with a peak in accuracy of 58% (*SEM* = 1.30) at 142 ms (Fig. S2f). Similar to Experiments 1 and 2, the cross-condition analysis revealed the generalisability of fearful face classifiers from supraliminal to subliminal conditions and vice versa, with peak in accuracies of 60% at around 145 ms (Fig. S2i & S2l)

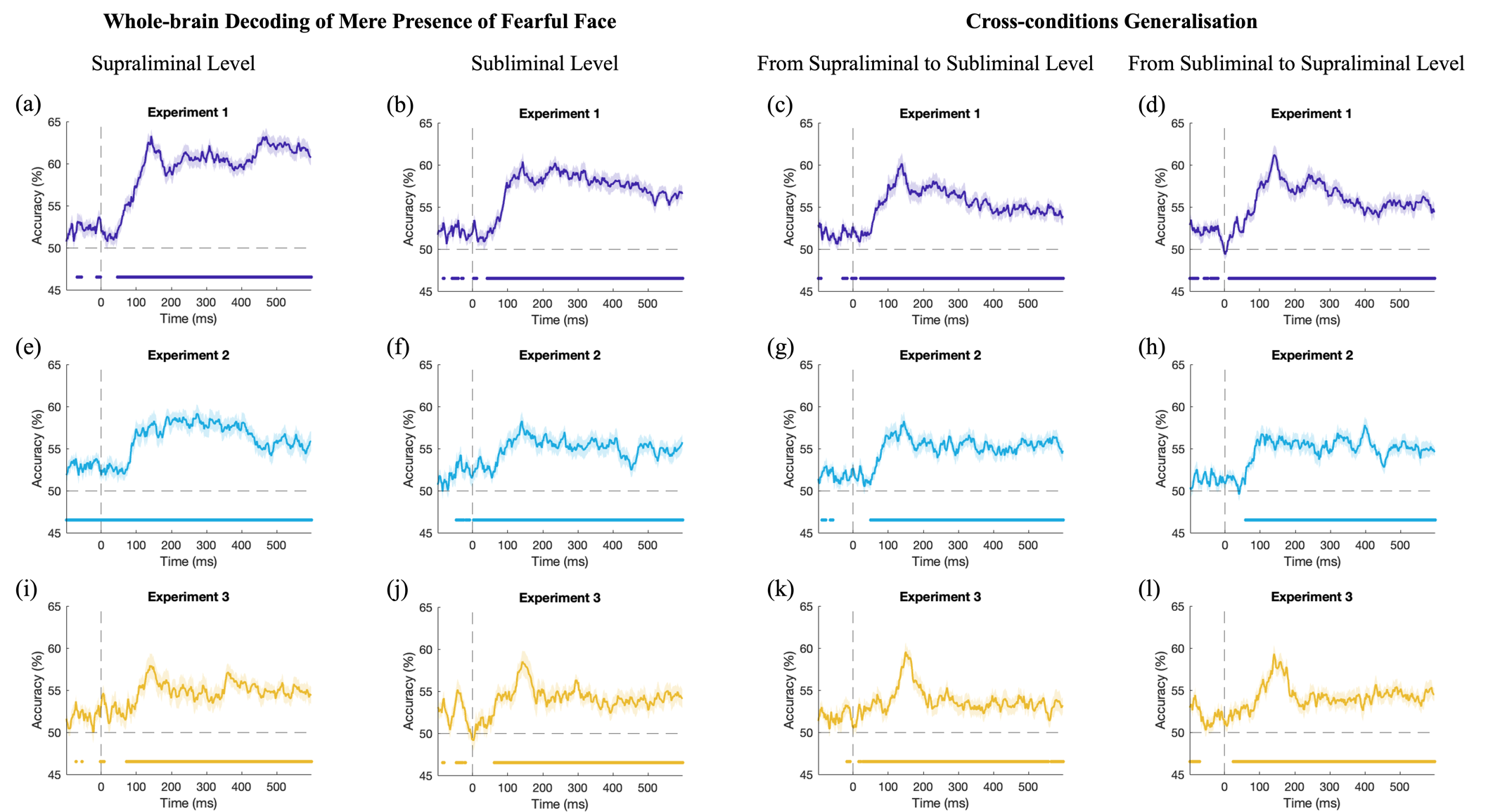

**Fig. S2.** Whole-brain Decoding and Cross-condition Decoding of Mere Presence of Fearful Face. Performance of fearful face decoding (presence vs. absence) using EEG signals across whole-brain electrodes at supraliminal level and subliminal level, and cross-condition decoding of fearful face from supraliminal to subliminal level and from subliminal to supraliminal level, for Experiment 1 (a-d), Experiment 2 (e-h) and Experiment (i-l) are shown. Time periods showing above-chance decoding (accuracy > 50%) are highlighted by the coloured dotted line in each plot, TFCE-corrected *p* < 0.05. The coloured shading in each plot represents standard error of the mean.

**Whole-brain decoding for the spatial location of fearful faces**

In Experiment 1, the spatial location (left vs. right) of fearful faces were decodable from whole-brain electrodes in the supraliminal conditions (Fig. S3a). Successful decoding was mainly found for time windows of 228-250 ms and 360-600 ms, with a peak at 56% (*SEM* = 1.22) at around 425 ms. Decoding of spatial location was not significantly above chance in the subliminal conditions (Fig. S3b).

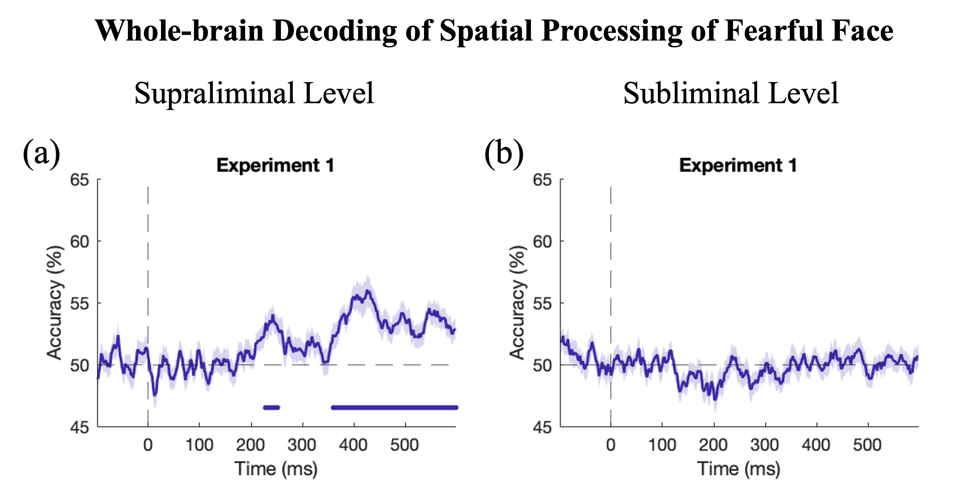

**Fig. S3.** Whole-brain Decoding of Spatial Processing of Fearful Face in Experiment 1. Decoding performance of spatial processing of fearful face (a) at supraliminal level and (b) subliminal level using whole-brain electrodes are shown. Time periods showing above-chance decoding (accuracy > 50%) are highlighted by the coloured dotted line in each plot, TFCE-corrected *p* < 0.05. The coloured shading in each plot represents standard error of the mean.
